# Drug degradation caused by *mce3R* mutations confers contezolid (MRX-I) resistance in *Mycobacterium tuberculosis*

**DOI:** 10.1101/2022.07.25.501497

**Authors:** Rui Pi, Xiaomin Chen, Jian Meng, Qingyun Liu, Yiwang Chen, Cheng Bei, Chuan Wang, Qian Gao

**Affiliations:** Key Laboratory of Medical Molecular Virology (MOE/NHC/CAMS), School of Basic Medical Sciences, Shanghai Medical College, Shanghai Institute of Infectious Disease and Biosecurity, Fudan University, Shanghai, China; Central Laboratory, Zhejiang Chinese Medicine and Western Medicine Integrated Hospital (Hangzhou Red Cross Hospital), Hangzhou, Zhejiang Province China; Shanghai Institute of Materia Medica, Chinese Academy of Sciences, Shanghai, China; National Clinical Research Center for Infectious Diseases, Shenzhen Third People’s Hospital, Shenzhen, Guangdong Province, China

**Author notes:** **Corresponding author**: Qian Gao, PhD, Key Laboratory of Medical Molecular Virology, School of Basic Medical Sciences, Fudan University, Room 201, Fuxing Building, Dongan Road No. 131, 200032, Shanghai, China.

**Keywords:** *Mycobacterium tuberculosis*, drug resistance, contezolid, Mce3R, Rv1936, degradation

## Abstract

Contezolid (MRX-I), a safer antibiotic of the linezolid oxazolidinone class, is a promising new antibiotic with potent activity against *Mycobacterium tuberculosis* (MTB) both *in vitro* and *in vivo*. To identify resistance mechanisms of contezolid in MTB, we isolated several *in vitro* spontaneous contezolid-resistant MTB mutants, which exhibited 16-fold increase in MICs of contezolid compared with the parent strain but was still unexpectedly susceptible to linezolid. Whole-genome sequencing revealed that most of the contezolid-resistant mutants bore mutations in the *mce3R* gene which encode a transcriptional repressor. The mutations in *mce3R* led to markedly increased expression of a monooxygenase encoding gene *Rv1936*. We then characterized Rv1936 as a putative flavin-dependent monooxygenase that catalyzes the degradation of contezolid into its inactive DHPO ring-opened metabolites, thereby conferring drug resistance. While contezolid is an attractive drug candidate with potent antimycobacterial activity and low toxicity, the occurrence of mutations in Mce3R should be considered when designing combination therapy using contezolid for treating tuberculosis.

**IMPORTANCE:** Tuberculosis (TB) is one of the leading causes of global death and the second deadliest infectious killer after COVID-19. Compared to drug-sensitive TB, the treatment of multidrug-resistant (MDR) and extensively drug-resistant (XDR) TB is more difficult and less effective due to longer regimens and higher potential for clinical adverse events. Despite the undisputed medical success of linezolid on MDR/XDR-TB therapy, this drug suffers from severe safety limitation. The new NMPA-approved drug contezolid, as an analogue of linezolid, exhibits a superior safety profile and potent antitubercular activity. Since the less-toxic contezolid is a promising drug candidate to optimize the current longer-duration MDR/XDR-TB therapy, it would be of significance to determine the resistance profiles of contezolid in MTB. Here, we present the first exploration of the frequency, mutational targets and molecular mechanisms of contezolid resistance in MTB, which could provide theoretical guidance for its future clinical application.

## INTRODUCTION

Tuberculosis (TB), caused by *Mycobacterium tuberculosis* (MTB), afflicts about 10 million patients and causes approximately 1.5 million deaths every year (1). Although the global treatment success rates is 86% for drug-susceptible TB, it decreases to 59% for multidrug-resistant (MDR) (1), 47% for pre-extensively drug-resistant (pre-XDR) (2) and 39% for XDR TB cases (3). The introduction of the oxazolidinone linezolid into tuberculosis treatment regimens has effectively improved treatment success rates for MDR/XDR MTB strains (4–6), but around 48% to 87% of patients on prolonged linezolid therapy experience adverse toxic effects mainly including reversible hematologic problems and irreversible neuropathies (4–9). To address this toxicity, several new oxazolidinone agents have been rationally designed to reduce the myelosuppression and monoamine oxidase inhibition (MAOI) that are associated with linezolid toxicities (10).

Contezolid (MRX-I) is a novel potent oxazolidinone developed by Shanghai MicuRx Pharmaceutical Co., Ltd., and was recently approved by the National Medical Products Administration (NMPA) of China for the treatment of complicated skin and soft tissue infections (cSSTI) (11–13). Nonclinical and clinical studies have confirmed that, compared with linezolid, contezolid exhibits significantly reduced myelosuppression and MAOI inhibition meanwhile maintaining equivalent or even higher antibacterial activity both *in vitro* and *in vivo* (10, 14–20). Contezolid also has excellent antitubercular activity against both susceptible and resistant MTB including MDR/XDR strains (MIC_50_=0.5 μg/ml, MIC_90_=1 μg/ml) (21), and its lower toxicity might allow contezolid to supplant linezolid in the long-term treatment regimens required to cure MDR/XDR-TB. However, to fully appreciate the potential of contezolid as an anti-TB agent, it would be advantageous to understand the frequency, location and molecular mechanisms of mutations associated with contezolid resistance in MTB.

To understand the development of contezolid resistance in MTB, the present study *in vitro* selected contezolid-resistant mutants, determined their resistance profiles, identified the mutational targets and explored the underlying mechanisms conferring the resistance.

## MATERIALS AND METHODS

### Bacterial cultures

*Escherichia coli* TOP10 and BL21 Star (DE3) strains used for cloning were grown in LB with or without 1.5% agar. MTB strains were grown in Middlebrook 7H9 broth supplemented with 0.2% (vol/vol) glycerol, 0.05% (vol/vol) tyloxapol, and 10% (vol/vol) oleic acid-albumin-dextrose-catalase (OADC) or on Middlebrook 7H10 agar plates supplemented with 0.5% (vol/vol) glycerol and 10% (vol/vol) OADC. All bacteria were grown aerobically at 37°C with constant agitation. When indicated, kanamycin and hygromycin was added at 25 μg/ml and 50 μg/ml respectively. contezolid was used for mutant selection and MIC determination at concentrations ranging from 2.5 to 64 μg/ml.

### Spontaneous contezolid-resistant mutant selection

For each ten independent log-phase cultures of laboratory reference strain H37Rv, 3 ml aliquots were concentrated and spread onto three 7H10-OADC plates containing 5 μg/ml contezolid (Shanghai MicuRx Pharmaceutical Co. Ltd.), corresponding to 5 times the minimum inhibitory concentration (MIC) of 1 μg/ml contezolid for H37Rv. The colony forming units (CFU) were calculated by serially diluting the original cultures 10^−4^- to 10^−5^-fold and then triplicate plating aliquots onto antibiotic-free 7H10-OADC plates. After incubation at 37°C for 4 weeks, colonies appearing on the contezolid containing plates were replated onto 7H10-OADC plates containing 2.5 μg/ml contezolid to confirm their resistance. The frequency of spontaneous contezolid resistance was estimated as the ratio between the number of resistant colonies obtained and the total CFU plated.

### Whole genome sequencing and data analysis

Genomic DNA of 108 contezolid-resistant colonies, and the parental H37Rv, was extracted with the cetyltrimethylammonium bromide-lysozyme method and used for the Illumina construction of 300 bp fragment length genomic libraries for paired-end sequencing on an Illumina HiSeq 2500 instrument. A previously validated pipeline was used for mapping the sequencing reads to the reference genome (22). In brief, the Sickle tool was used for trimming and sequencing reads with Phred base quality scores above 20 and lengths longer than 30 bp were kept for analysis. Mutations were called with at least 2 reads in both forward and reverse directions, using MTB H37Rv strain (NC_000962.2) as a reference. Sequencing reads were mapped to the reference genome using Bowtie 2 (version 2.2.9) and SAMtools (v1.3.1) was used for SNP calling with mapping quality greater than 30. Fixed mutations (frequency ≥ 75%) were identified using VarScan (v2.3.9) with at least 10 supporting reads with the strand bias filter option on. SNPs in repetitive regions of the genome (e.g., PPE/PE-PGRS family genes, phage sequences, insertion, or mobile genetic elements) were excluded. Small INDELs were identified by VarScan (v2.3.9) using the mpileup2indel function.

### MIC determination

MIC values were determined in Middlebrook 7H9-OADC broth in 96-well plates using a standard microdilution method. Serial two-fold dilutions of antibiotics were prepared in 96-well plates. Early-log phase (OD_600_=0.5-1.0) MTB cultures were diluted to ~ 1 × 10^5^ CFU/ml (OD ≈ 0.001) and then equal volumes (100 μl) of diluted cultures were added to each well to yield a final testing volume of 200 μl. The plates were incubated at 37 °C for 2-3 weeks and then MICs were determined visually as the minimum drug concentration preventing bacterial growth. Contezolid and linezolid were generously provided by Shanghai MicuRx Pharmaceutical Co. Ltd. and solubilized in dimethylsulfoxide (DMSO) as a stock solution of 5 mg/ml. For drug susceptibility determination, these stocks were diluted in 7H9-OADC broth to final concentrations of 64, 32, 16, 8, 4, 2, 1 and 0.5 μg/ml. To exclude the bacteriostatic effect of the solvent, DMSO was diluted as well for drug-free control.

### Plasmid construction and transformation

The high-copy-number non-integrative plasmid pSMT3-LxEGFP was used for the genetic overexpression and complementation. Briefly, the gene containing DNA fragments were PCR-amplified from MTB H37Rv with the primers shown in Table S1, and then inserted into plasmid pSMT3-LxEGFP restricted with BamHI and HindIII (Thermo Scientific) using T4 DNA ligase. The recombinant plasmids containing *Rv1936, Rv1938* genes and the *Rv1936-Rv1941* operon were then independently electroporated into the transformation competent wild type (WT) H37Rv. As for *mce3R* mutant complementation, the recombinant plasmid containing a WT copy of *mce3R* was electroporated into the transformation competent *mce3R*_417_527del mutant, which has an 111 basepair (bp) deletion in *mce3R*.

### RNA extraction and quantitative PCR (qPCR)

Cultures were grown to early-log phase (OD_600_ = 0.5-1.0) and harvested at 4°C. Cell pellets were re-suspended in 1 ml TRIzol reagent (Invitrogen) and subjected to bead-beating with 0.1 mm silica beads at maximum speed for 30 seconds three times, with cooling on ice between shaking. After centrifugation at 13200 rpm for 1 minute at 4°C, the supernatant was transferred to a phase lock gel tube (TIANGEN) containing 400 μl chloroform, inverted vigorously for 30 seconds and centrifuged at 13200 rpm for 15 minutes. The upper aqueous phase (about 500 μl) was then transferred to a new tube and the RNA precipitated with 450 μl isopropanol for 30 minutes at room temperature. RNA was washed sequentially with 80% and pure ethanol, and the pellet then dissolved in 50 μl RNAase free water. The total RNA yield was quantified with a Nanodrop (Thermo Scientific). cDNA was synthesized using the PrimeScript™ RT reagent Kit (TAKARA) following the manufacturer’s instructions. qPCR reactions were performed with iQ SYBR green supermixture kit (TAKARA) in the CFX96TM Real-Time PCR System (Bio-Rad). The sequences of all primers are listed in Table S1.

### Co-incubation sample collection and metabolites profiling

The contezolid stock solution was added to 4 ml of stationary phase H37Rv cultures at final concentrations of 2μM and 10μM (final DSMO concentrations 0.025% and 0.125%) for susceptible and resistant strains, respectively. The mixtures were incubated at 37 °C for 15 hours, heated to 80°C for 30 minutes to kill the MTB and then subjected to bead-beating with 0.1 mm silica beads at maximum speed three times for 30 seconds each with cooling on ice between shakes. After centrifugation at 10000 rpm for 3 minutes at 4°C, the supernatants were directly analyzed by ultraperformance liquid chromatography (UPLC)/triple time-of-flight mass spectrometry (TOF MS) to identify contezolid and its metabolites as previously described (23). In brief, 10 μl of supernatant was injected into the Acquity UPLC HSS T3 column (100 × 2.1 mm i.d., 1.8 μm) eluted with a linear gradient. The mobile phase consisting of 0.05% formic acid in water and methanol with the flow rate maintained at 0.45 ml/min. Contezolid and its metabolites were detected using a triple TOF 5600+ MS/MS system (AB Sciex, Concord, Ontario, Canada) in the positive electrospray ionization mode, with the mass range set at *m/z* 100 – 1000. Information-dependent acquisition (IDA) was used to trigger the acquisition of MS/MS spectra for contezolid and its metabolites. A real-time multiple mass defect filter was used for the IDA criteria.

### Quantification and statistical analysis

Graphics and data analyses were performed in Prism version 7.0 software (GraphPad) and SnapGene version 4.3.6 software (GSL Biotech LLC).

### Accession codes

The whole genome sequencing (WGS) data were deposited in the Genome Sequence Archive (https://bigd.big.ac.cn/gsa) under accession number CRA007448.

## RESULTS

### *In vitro* selection of contezolid-resistant MTB mutants

Ten independent cultures of the pan-susceptible WT H37Rv strain were grown to log phase and cells from each 3 ml culture were spread onto 7H10-OADC plates containing 5 μg/ml contezolid, five-fold higher than the MIC of 1 μg/ml for the WT strain (Table 1). From the total of about 2.1×10^10^ CFU plated, there arose almost 6000 resistant colonies, yielding a spontaneous contezolid resistance frequency of approximately 3 × 10^−7^.

**TABLE 1.** Mutations identified in 106 contezolid-resistant strains of *M. tuberculosis*

| Mutation type | Number of strains | Percentage |
| --- | --- | --- |
| SNPs in *mce3R* | 59 | 55.6% |
| Nonsynonymous mutations | 47 | 44.3% |
| Nonsense mutations | 12 | 11.3% |
| Indels in *mce3R* | 40 | 37.8% |
| Insertion mutations | 11 | 10.4% |
| Deletion mutations | 27 | 25.5% |
| Multiple variants in *mce3R* | 2 | 1.9% |
| Unknown | 7 | 6.6% |
| Total | 106 | 100% |

### WGS identified *mce3R* as the most frequently mutated gene

A total of 120 resistant colonies, representing equally the 10 original cultures, were replated onto 7H10 agar plates containing 2.5 μg/ml contezolid and confirmed to be resistant. The genomic DNA was successfully extracted from 108 of the 120 resistant isolates and subjected to WGS. Two isolates with sequencing depths less than 15 were excluded from further analysis. The WGS data revealed that 93.4% (99/106) of the resistant isolates had mutations in *mce3R* (*Rv1963c*), which encodes a TetR family transcriptional regulator, while the remaining 6.6% (7/106) harbored no confirmable mutations. The mutations in *mce3R* were scattered throughout the entire gene with no mutagenic “hot-spot” regions (Fig. 1A). A total of 35 nonsynonymous mutations were found in 47 resistant strains and 6 nonsense mutations were found in 12 strains. In addition, 10 different insertions were present in 11 resistant strains and 14 different deletions were found in 27 resistant strains (Table 1). Although most insertions and deletions involved a single nucleotide causing a frameshift, there were 5 resistant strains that contained either a 98bp (417_527del) or a 111bp (583_680del) deletion within the *mce3R* gene. Additionally, in two resistant strains the WGS data showed multiple mutations within the 339-466 base region of the *mce3R* gene, which were confirmed by Sanger sequencing (Fig. 1B).

**FIG 1.**
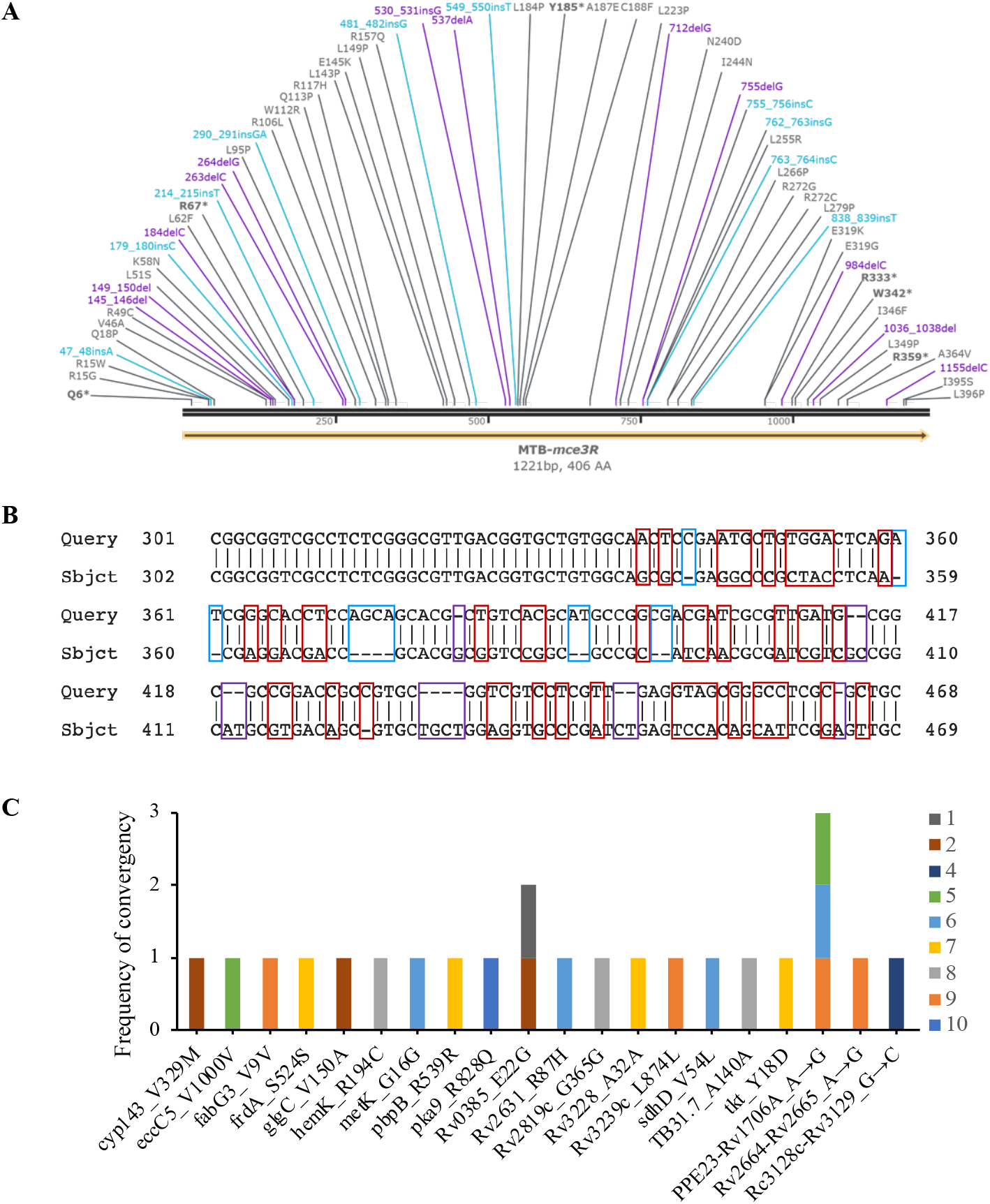
The genetic variants in contezolid-resistant *M. tuberculosis*. (A) Distribution of *mce3R* mutations associated with contezolid resistance. The full length of *mce3R* gene is 1221bp, encoding a putative protein of 406 amino acids. SNPs are showed in gray with nonsense mutations in bold, with numbers that correspond to the amino acid positions. Insertion and deletion mutations are shown in indigo and purple, respectively, with numbers indicating base positions. (B) Two contezolid-resistant strains harbored multiple mutations within the 339-446 base region of the *mce3R* gene. SNPs, insertions and deletions are indicated by red, blue and purple boxes, respectively. The Query and Sbjct sequences belong to contezolid-resistant strains and the wild-type H37Rv strain, respectively. The numbers indicate the base position of the *mce3R* gene. (C) Distribution of hitchhiking mutations identified in contezolid-resistant strains by WGS analysis, among 10 independent selection cultures indicated with different colors.

By exploring the entire genomes of the 106 contezolid-resistant mutants, we also identified twenty other mutations in 17 isolates, but 90% of these mutations (18/20) were found in strains that also contained *mce3R* mutations. Twelve of the 20 other mutations were intergenic or synonymous and predicted to have no impact on the function of their encoded proteins, and most (18/20) mutations were found in a resistant strain obtained from just one of the ten original cultures used for mutant selection. Therefore, it was presumed that these 20 mutations fixed in MTB were forms of genetic hitchhiking unrelated to contezolid resistance.

### Determination of contezolid resistance target and its cross-resistance with linezolid

To confirm that the *mce3R* mutations were responsible for the contezolid resistance, we cloned the entire WT *mce3R* gene into high-copy-number non-integrative plasmid pSMT3-LxEGFP and complemented one of the *mce3R* mutants with a large *mce3R* deletion (417_527del) using the constructed plasmid. The complemented *mce3R* deletion strain recovered its susceptibility to contezolid with the same MIC of 1 μg/ml as the WT H37Rv parent strain (Table 2), proving that the *mce3R* mutations were responsible for contezolid resistance.

**TABLE 2.** MIC values for contezolid and linezolid in MTB strains analyzed in this study.

| Strain ID | Characteristic | MIC (µg/ml) |  |
| --- | --- | --- | --- |
|  |  | contezolid | linezolid |
| MTB-260 | *mce3R*_R15G mutant | 16 | 1 |
| MTB-243 | *mce3R*_Q6stop mutant | 16 | 1 |
| MTB-224 | *mce3R*_838_839insT mutant | 16 | 1 |
| MTB-197 | *mce3R*_264delG mutant | 16 | 1 |
| MTB-21 | *rrl*_G2270T mutant | 16 | 16 |
| MTB-33 | *rrl*_G2814T mutant | 16 | 32 |
| MTB-12 | *rplC*_T460C mutant | 16 | 16 |
| H37Rv | WT strain | 1 | 1 |
| $\Delta mce3R$ | *mce3R*_417_527del mutant | 16 | 1 |
| $\Delta mce3R$ O/E *mce3R* | *mce3R*_417_527del mutant complemented with a WT copy of *mce3R* | 0.5-1 | 1 |
| H37Rv O/E *Rv1936* | MTB overexpressing *Rv1936* | 16 | 1 |
| H37Rv O/E *Rv1938* | MTB overexpressing *Rv1938* | 1 | 1 |
| H37Rv O/E *Rv1936-Rv1941* | MTB overexpressing *Rv1936-Rv1941* operon | 16 | 1 |

We then evaluated the effect of the *mce3R* mutations on both contezolid and linezolid resistance. The MTB strains harboring different *mce3R* mutations (R15G, Q6stop, 838_839insT, 264delG and 417_527del) uniformly exhibited a 16-fold increase in contezolid resistance (MIC=16 μg/ml), but remained susceptible to linezolid (MIC=1 μg/ml) (Table 2). In contrast, our previously isolated linezolid-resistant MTB strains, harboring canonical oxazolidinone resistance mutations in *rrl* or *rplC* (24), exhibited cross-resistance to contezolid, with MICs (16 μg/ml) equal to that of the *mce3R* mutants (Table 2). These results suggested that mutations in the *mce3R* gene confer resistance specifically to contezolid and do not constitute a generalized oxazolidinone resistance mechanism.

### Overexpression of Rv1936 induced by Mce3R mutations confers contezolid resistance

As Mce3R is a TetR transcriptional repressor that negatively regulated genes involved in lipid metabolism, redox reactions and stress resistance (25–27), we reasoned that genes in its regulon might be involved in the molecular mechanism of contezolid resistance in MTB. Previous proteomic results directed our attention to two genes located in the *Rv1936-Rv1941* operon which was negatively regulated by Mce3R, in that their coding products, namely a monooxygenase encoded by *Rv1936* and an epoxide hydrolase EphB encoded by *Rv1938*, have been found to be dramatically overabundant in the *mce3R* knock-out mutant than in the WT H37Rv strain (25). Similar enzymes have been shown to be generally involved in the detoxification and catabolism of xenobiotics in microorganisms (28, 29). To better understand their association with contezolid resistance, we constructed strains containing plasmids overexpressing *Rv1936, Rv1938* or the *Rv1936-Rv1941* operon. Real-time fluorescent quantitative PCR (RT-qPCR) confirmed that *Rv1936* and *Rv1938* genes were markedly co-upregulated in our *mce3R* mutants and the strain containing plasmid with the *Rv1936*-*Rv1941* operon, as well as individually upregulated in their corresponding overexpressing strains, whereas were expressed at the WT level in the *mce3R* mutant complemented with a WT copy of *mce3R* (Fig. 2). We then tested their contezolid susceptibilities and found that MTB strains containing plasmids overexpressing *Rv1936* or the *Rv1936-Rv1941* operon had a MIC of 16 μg/ml, showing a level of resistance as high as that displayed by the *mce3R* mutants, whereas the strain only overexpressing *Rv1938* had a MIC of 1 μg/ml to contezolid, remaining unchanged susceptibility as with the WT strain (Table 2). These findings suggest that the mutations in *mce3R* abrogate its negative regulatory function, leading to overexpression of the monooxygenase encoded by *Rv1936*, which then confers contezolid resistance in MTB.

**FIG 2.**
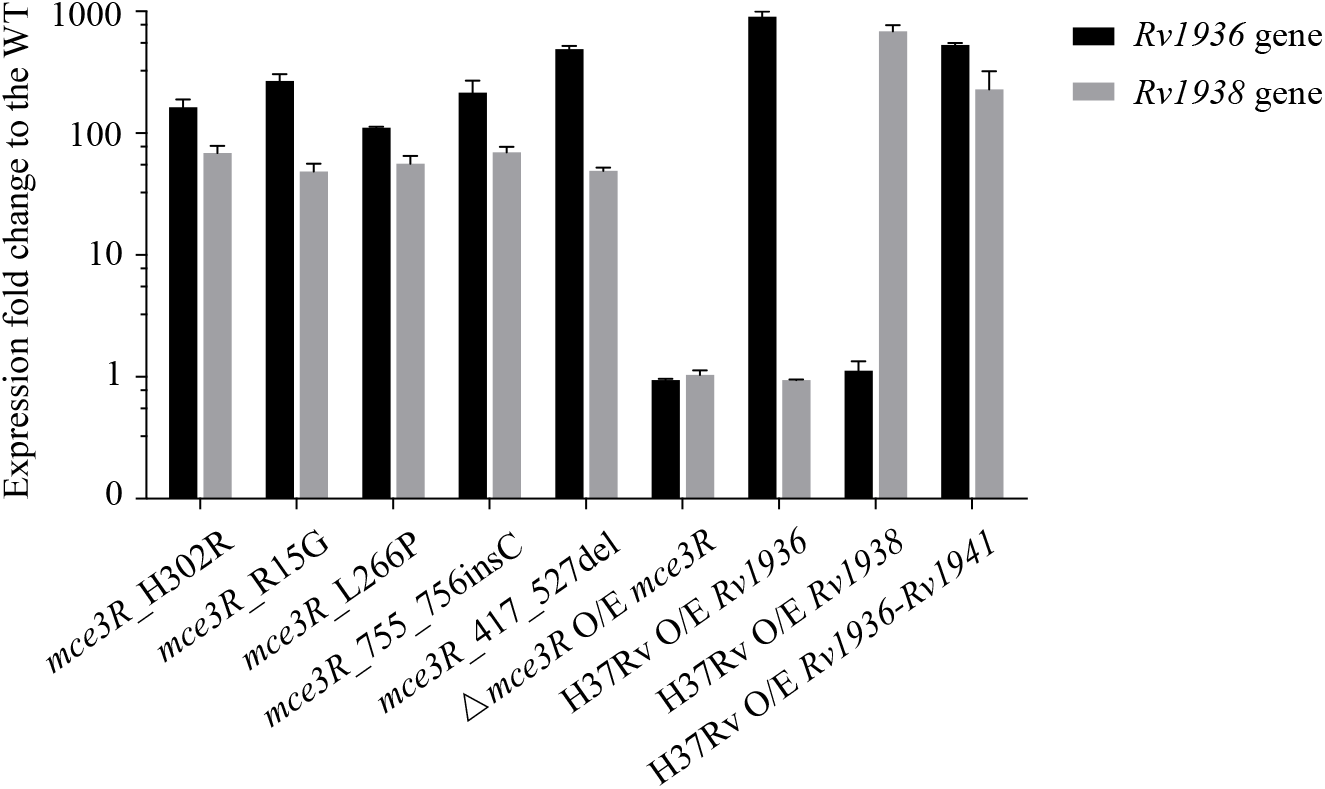
Gene transcript level detected by RT-qPCR. The figure shows the change in transcriptional expression, relative to the WT H37Rv, of the *Rv1936* and *Rv1938* genes in five *in vitro*-isolated *mce3R* mutants, the *mce3R*_417__521del mutant complemented with a WT copy of *mce3R* gene (Δ*mce3R* O/E *mce3R*), and three MTB strains with plasmids overexpressing (O/E) the indicated genes.

### Rv1936 degrades contezolid in resistant MTB stains

We next sought to relate the overexpression of Rv1936 to the mechanism of contezolid resistance in MTB. As Rv1936 is non-essential for the *in vitro* growth of MTB (30), which was unlikely to be the target of contezolid, we hypothesized that Rv1936 inactivate contezolid through drug degradation or modification. Sequence alignment analysis found that Rv1936 shares ~ 98% sequence homology with the flavin-dependent oxidoreductase MelF in *Mycobacterium marinum* (Fig. 3A), indicating a reliable enzymatic activity of this MTB monooxgenase (31). Moreover, given that contezolid in human is mainly metabolized by the liver enzyme flavin-containing monooxygenases 5 (FMO5) (23), we reasoned that Rv1936 might act as a putative FMO that degrades contezolid to confer resistance in MTB.

**FIG 3.**
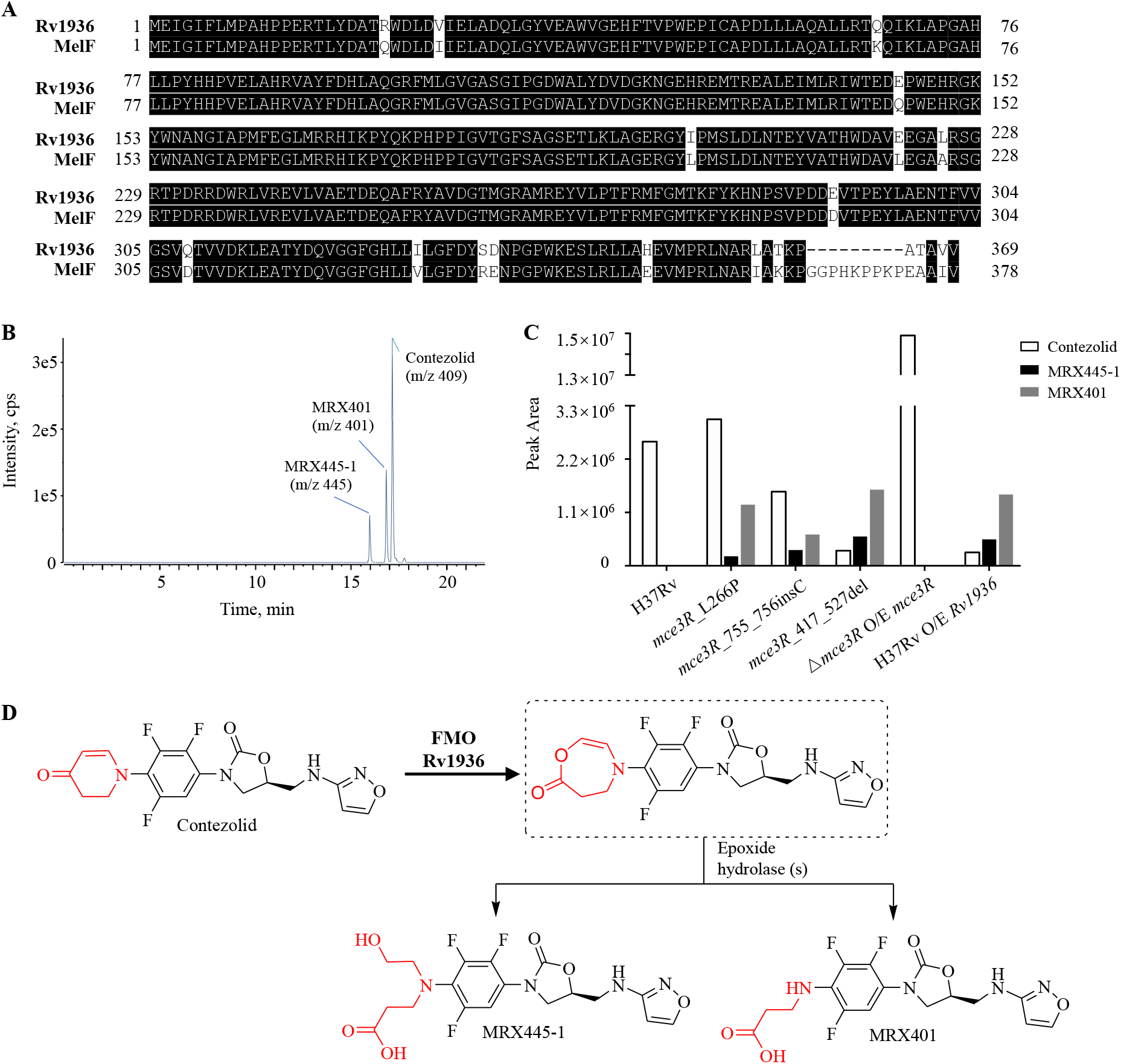
MTB Rv1936-promoted contezolid degradation. (A) Alignment of the amino acid sequences of the MTB Rv1936 and the flavin-dependent oxidoreductase MelF in *M. marinum*. Highlighted amino acids represent identity between sequences. The numbers indicate the amino acid position of Rv1936 or MelF. (B) Chromatogram representation of the parent contezolid and its two principal degradation products MRX445-1 and MRX401detected by UPLC/triple TOF MS, with their corresponding m/z values indicated. The retention time and mass are confirmed as described before (23). (C) Comparative quantification of contezolid, MRX445-1 and MRX401 based on their chromatographic peak area in the WT H37Rv, in the three contezolid-resistant *mce3R* mutants (L266P, 755_756insC and 417_527del), in Δ*mce3R* complemented with a constitutive expression vector for *mce3R* (Δ*mce3R* O/E *mce3R*) and in the WT H37Rv that constitutively expresses *Rv1936* (H37Rv O/E *Rv1936*). All strains were treated with 10 μl contezolid for 15 hours, with the exception of the WT H37Rv being treated with 2 μl contezolid to avoid drug inhibition effect. (D) Schematic representation of oxidation of contezolid by Rv1936, and subsequent hydrolysis of its DHPO ring-expanding enol lactone, the oxidative intermediate product indicated in dotted box, into two principal DHPO ring-opened metabolites MRX445-1 and MRX401. Chemical groups indicating the key DHPO ring-opening process of contezolid degradation are highlighted in red. The structures of contezolid and MRX445-1 were confirmed with authentic molecules, and the proposed structure of MRX401 is consistent with available high-resolution mass spectrometry data and its biotransformation pathway.

To test the Rv1936-promoted contezolid degradation hypothesis, we grew MTB cells for 15 hours in the presence of contezolid and used UPLC/Triple TOP MS to identify the parent drug as well as its metabolites in the coincubation mixtures. Overall, the most abundant components were the parent drug contezolid (C_18_H_15_N_4_O_4_F_3_) and its two 2,3-dihydropyridin-4-one (DHPO) ring-opened metabolites: MRX445-1 (C_18_H_19_N_4_O_6_F_3_) and MRX401 (C_16_H_15_N_4_O_5_F_3_) (Fig. 3B, 3C). In detail, compared with results obtained from contezolid incubated with contezolid-susceptible MTB strains (i.e. WT H37Rv and the *mce3R* complemented mutant strain), we observed a dramatic reduction on the amount of the parent drug contezolid in the incubation samples of contezolid treated with contezolid-resistant MTB strains (i.e. *mce3R*_755_756insC mutant, *mce3R*_417_527del mutant, *mce3R*_L266P mutant and the strain overexpressing the *Rv1936* gene) (Fig. 3C). Meanwhile, the amount of two DHPO ring-opened metabolites MRX445-1 and MRX401, as the consequence of contezolid degradation, increased in the resistant mixtures relative to the susceptible ones (Fig. 3C). Our findings suggested that the overexpression of the putative FMO Rv1936 promote the degradation of contezolid, presumably by oxidizing the DHPO ring-expanding process for subsequent hydrolytic degradation (Fig. 3D), thereby conferring drug resistance in MTB.

## DISCUSSION

Contezolid is a new oxazolidinone antibiotic that retains the potent antitubercular activity of linezolid but is less-toxic and could potentially replace linezolid in the treatment of drug-resistant TB (10, 18–21). This study sought to understand the development of resistance to contezolid by *in vitro* isolating and characterizing spontaneous contezolid-resistant MTB mutants. Surprisingly, 93.4% of the contezolid resistant strains had mutations in *mce3R*, a TetR-like transcriptional repressor. A wide variety of mutations were found throughout the *mce3R* gene, suggesting that any loss-of-function mutations would confer resistance. Consistently, complementation of mutant with a WT copy of *mce3R* restored contezolid susceptibility. The inactivation of Mce3R in these mutant strains led to increased expression of *Rv1936*, which appeared to code for the critical functional enzyme that catalyzed contezolid for degradation, thereby reducing the level of contezolid and conferring resistance in MTB.

In humans, contezolid is principally metabolized into MRX445-1 and MRX459 via the Baeyer-Villiger oxidation of DHPO ring which is specifically catalyzed by the liver FMO5, followed by subsequent hydrolytic DHPO ring-opening process (23). In the current study, we have uncovered that MTB exploits the similar mechanism to counter the antitubercular activity of contezolid, by degrading the parent drug into its DHPO ring-opened metabolites MRX445-1 and MRX401. Based on the established literature of structure-activity relationship (SAR) data for oxazolidinones, metabolites of contezolid are not expected to exhibit potent antibacterial activity (18). Further MIC tests have confirmed that the two human metabolites MRX445-1 and MRX459 showed no antibacterial activities against Gram-positive pathogens including *Staphylococcus aureus (S. aureus), Enterococcus faecium, Enterococcus faecalis* and *Streptococcus pneumoniae* (32). Given that MRX401 is the dealkylated metabolite derived from MRX445-1, it should also exhibit no antibacterial activity.

Mutations in *mce3R*, or overexpression of *Rv1936* contributed to contezolid resistance in MTB, however, did not confer resistance to linezolid, because the chemical structure of linezolid lacks the DHPO ring targeted by the putative FMO Rv1936. Through searching on the NCBI BLAST (https://blast.ncbi.nlm.nih.gov/Blast.cgi) and the mycobrowser database (https://mycobrowser.epfl.ch/releases), we only found orthologues of the MTB Mce3R and Rv1936 in *Mycobacterium bovis*, *M. canettii*, *M. marinum*, *M. ulcerans* and *M. vanbaalenii*. Thus, it is possible that the degradation of contezolid as a mechanism of resistance may be limited in bacteria with a suitable monooxygenase whose expression is controlled by a non-essential negative regulator, such as *mce3R*. While the drug-degradation mechanism of contezolid resistance appears to be unique to a few mycobacteria, degradation of antibiotics is a common mechanism through which pathogens develop drug resistance. One of the most prominent examples is the enzymatic degradation of β-lactam antibiotics through the acquisition of β-lactamases that hydrolyze the β-lactam ring (33). Although MTB possesses strong, constitutive β-lactamase activity that renders it intrinsically resistant to β-lactam antibiotics (34–38), the current study provides another example of the acquired degradation mechanism conferring drug resistance in MTB.

Given that the contezolid resistance could be achieved by overexpressing *Rv1936* alone in the WT H37Rv, rather than overexpressing *Rv1938* alone, we reasoned that oxidation of the DHPO ring catalyzed by monooxygenase Rv1936 is pivotal for the degradation of contezolid, which appears to be the rate-limiting step, and MTB should have other intrinsically abundant epoxide hydrolase(s) rather than Rv1938 to catalyze the subsequent hydrolysis of the oxidized DHPO ring. Preliminary studies suggested that MTB genome encodes other six putative monooxygenases (i.e. Rv0892, Rv0565c, Rv3854c, Rv1393c, Rv3049c and Rv3083) capable of performing Baeyer-Villiger oxidations and seven putative epoxide hydrolases (i.e. Rv3617, Rv1938, Rv1124, Rv2214c, Rv3670, Rv0134 and Rv2740, corresponding to Eph A-G) (39, 40). Further enzymatic assays have confirmed the oxidating activity of Rv3854c, Rv3049c and Rv3083 (39, 41), as well as the hydrolyzing activity of Rv3617, Rv1938, Rv1124, Rv0134 and Rv2740 (42–45). In particular, Rv3854c (EthA) and Rv3083 (MymA) have been confirmed to similarly function as the activating enzymes of the anti-TB prodrug ethionamide, whereas individually activate different sulfur-containing compounds, suggesting the overlap in enzymatic function and substrate specificity of these monooxygenases in MTB (41). It remains for further study to identify whether these six monooxygenases in MTB mentioned above could catalyze the oxidation of DHPO ring and which epoxide hydrolase(s) participated in the subsequent hydrolytic ring-opening process.

Although contezolid has been used clinically for the treatment of cSSTI, very little is known about the potential mechanisms of its clinical resistance. The only published data on contezolid resistance found mutations in the canonical oxazolidinone gene targets *rrl* and *rplC* in spontaneous *in vitro*-isolated contezolid-resistant strains of *S. aureus* (46). In the present study, however, none of our 106 *in vitro*-isolated resistant MTB strains harbored mutations in *rrl* or *rplC*, even though we showed that MTB strains with mutations in these genes are resistant to contezolid. In subsequent work we isolated additional contezolid resistant strains and found mutations in *rrl* or *rplC*. The predominance of *mce3R* mutations in contezolid-resistant strains is likely because these inactivating mutations can occur anywhere in the gene and are found at an estimated frequency of approximately 3 × 10^−7^, which is nearly one log higher than the previously reported mutation frequencies in *rrl* and *rplC* of 2 × 10^−8^ ~ 5 × 10^−9^ (47, 48). Moreover, such mutational bias raises the possibility that mutations in the nonessential gene *mce3R* might display lower fitness cost on MTB strains than in the essential genes *rrl* and *rplC*. Interestingly, there remained seven *in vitro*-isolated contezolid-resistant MTB strains harboring neither mutations in the mechanistic *rrl* and *rplC* targets, nor in our newly identified non-mechanistic *mce3R* target, suggesting there may be other alternative mutational targets and mechanisms for contezolid resistance in MTB.

Another intriguing observation is that the contezolid resistance conferring mutations in *mce3R* were highly polymorphic and included 35 non-synonymous mutations, 6 nonsense mutations and 24 frameshift mutations scattered throughout the whole *mce3R* gene rather than in a “hot-spot” region. Similar polymorphic patterns have been observed for mutations in *pncA* that confer resistance to pyrazinamide and in *Rv0678* that confer resistance to bedaquiline. Studies based on *in silico* structure analysis illustrated that, in addition to frameshift mutations, mutations can inactivate the protein function by impairing protein conformational flexibility, folding, stability, or catalytic activity (49–51). Further investigations are warranted to elucidate the effect of the contezolid resistance conferring mutations on Mce3R protein function. Although the frequency of *mce3R* mutations, 3 × 10^−7^ is relatively high, this may not discourage the use of contezolid against tuberculosis as the two critical first-line anti-TB drugs isoniazid and ethambutol have comparable even higher resistance frequencies of 1.78 × 10^−6^ and 1 × 10^−7^, respectively (52, 53). MTB resistance occurs rapidly if any single anti-TB drug is used in monotherapy, however, the clinical successful induction and maintenance therapy for TB is always based on strategy with multidrug therapy approach which could minimize the risk of acquiring drug resistance and achieve the maximum therapeutic efficacy. In addition, our study was based on *in vitro* selection of resistant isolates derived from the WT H37Rv strain, whereas the mutational location and frequency of *in vivo*-developed contezolid resistance in clinical MTB strains may be different.

In conclusion, this study identified the *mce3R* gene as a novel target for mutations conferring resistance to contezolid but not linezolid in MTB. The mutations in *mce3R* lead to upregulated expression of Rv1936, a putative FMO that initiates the degradation of contezolid into inactive metabolites. Our results provide fundamental insight into the *in vitro*-raised contezolid resistance in MTB, and suggest that the occurrence of mutations in *mce3R* gene should be considered when designing combination therapy using contezolid for treating tuberculosis.

## ACKNOWLEDGMENTS

This study was supported by the National Natural Science Foundation of China (81661128043, 81871625 to QG) and Shanghai Municipal Science and Technology Major Project (2018ZX10715-012).

We thank Howard E, Takiff and Qinglan Wang for critically reading and editing the manuscript, and Shanghai MicuRx Pharmaceutical Co. Ltd. for providing the two oxazolidinone antibiotics, contezolid (MRX-I) and linezolid.

The authors declare no competing interests.

